# Comparative genomics of *Alternaria* species provides insights into the pathogenic lifestyle of *Alternaria brassicae* – a pathogen of the *Brassicaceae* family

**DOI:** 10.1101/693929

**Authors:** Sivasubramanian Rajarammohan, Kumar Paritosh, Deepak Pental, Jagreet Kaur

## Abstract

*Alternaria brassicae*, a necrotrophic pathogen, causes *Alternaria* Leaf Spot, one of the economically important diseases of *Brassica* crops. Many other *Alternaria spp.* such as *A. brassicicola* and *A. alternata* are known to cause secondary infections in the *A. brassicae*-infected Brassicas. The genome architecture, pathogenicity factors, and determinants of host-specificity of *A. brassicae* are unknown. In this study, we annotated and characterised the recently announced genome assembly of *A. brassicae* and compared it with other *Alternaria spp.* to gain insights into its pathogenic lifestyle. Additionally, we sequenced the genomes of two *A. alternata* isolates that were co-infecting *B. juncea*. Genome alignments within the *Alternaria spp.* revealed high levels of synteny between most chromosomes with some intrachromosomal rearrangements. We show for the first time that the genome of *A. brassicae*, a large-spored *Alternaria* species, contains a dispensable chromosome. We identified 460 *A. brassicae*-specific genes, which included many secreted proteins and effectors. Furthermore, we have identified the gene clusters responsible for the production of Destruxin-B, a known pathogenicity factor of *A. brassicae*. The study provides a perspective into the unique and shared repertoire of genes within the *Alternaria* genus and identifies genes that could be contributing to the pathogenic lifestyle of *A. brassicae*.

## Introduction

The genus *Alternaria* belonging to the class of *Dothideomycetes* contains many important plant pathogens. Diseases in the *Brassicaceae* family caused by *Alternaria spp.* result in significant yield losses^1^. *Alternaria spp.* have a wide host range within the *Brassicaceae*, infecting both the vegetable as well as the oilseed crops. Some of the most damaging species include *Alternaria brassicae, A. brassicicola, A. alternata, A. raphani, A. japonicus*, and *A. tenuissima. A. brassicae* preferentially infects the oleiferous *Brassicas* while the others are more devastating on the vegetable *Brassicas. A. brassicae* is particularly more damaging in the hilly regions of the Indian subcontinent, where conducive climatic conditions allow it to profusely reproduce and cause infections on almost all parts of the plant. Extensive screening for resistance to *A. brassicae* in the cultivated *Brassica* germplasms has not revealed any source of resistance^2^.

The factors that contribute to the pathogenicity of *A. brassicae* are relatively unknown. Pathogenicity of many *Alternaria spp.* has been mainly attributed to the secretion of host-specific toxins (HSTs). HSTs induce pathogenesis on a rather narrow species range and are mostly indispensable for pathogenicity. At least 12 *A. alternata* pathotypes have been reported to produce HSTs and thereby cause disease on different species^3^. Many of the HST producing genes/gene clusters have been found on supernumerary chromosomes or dispensable chromosomes^4^. *A. brassicae* has been reported to produce low molecular weight cyclic depsipeptides named destruxins. Destruxin B is known to be a major phytotoxin and is reported to be a probable HST of *A. brassicae*^*5*^. Additionally, a proteinaceous HST (ABR-toxin), was isolated from the spore germination fluid of *A. brassicae* but was only partially characterised^6,7^.

Genome sequencing and comparative analysis can help identify shared and species-specific pathogenicity factors in closely-related species. Genomic information for nearly 26 *Alternaria spp.* including *A. brassicae* is currently available and has contributed immensely to clarify the taxonomy of the *Alternaria* genus^8^. However, comparative analyses to identify pathogenicity factors that confer the ability to infect a wide range of hosts have not been carried out. Most of the genomic information available for *Alternaria spp.* has been generated by shotgun sequencing approaches and hence is fragmented. A contiguous genome assembly is essential, especially when the aim is to identify and characterise pathogenicity factors or effectors, which are often present in rapidly evolving repeat-rich regions of the genome^9^. Additionally, contiguous genome assemblies enable an accurate prediction of genes and gene clusters that are involved in various secondary metabolic processes, many of which are implicated to have an important role in pathogenicity. Long reads generated from Pacific Biosciences (PacBio) single-molecule real-time (SMRT) sequencing technology and Oxford Nanopore sequencing technology enable the generation of high-quality genome assemblies at affordable costs. Besides the recently announced near-complete genome sequence of *A. brassicae*^10^, three other near-complete genomes of *Alternaria spp.* have been reported recently^11-13^.

*Alternaria* Leaf spot in the field usually occurs as a mixed infection of *A. brassicae* and other *Alternaria* species, such as *A. brassicicola* and *A. alternata*. It is however not known whether the *A. alternata* infecting *Brassicas* represent a separate pathotype with a different range of host-specific toxin(s) or are just opportunistic pathogens. We, therefore, carried out Nanopore-based sequencing of two *A. alternata* isolates that were recovered from an *A. brassicae*-infected *B. juncea* plant.

Given the invasiveness of *A. brassicae* and the lack of information on its pathogenicity factors, we undertook the current study to 1) functionally annotate and characterise the recently announced genome of *A. brassicae*, 2) sequence and analyse the genomes of two *A. alternata* isolates co-infecting *B. juncea* with respect to the genome of *A. alternata* isolated from very divergent hosts, 3) analyse the repertoire of CAZymes, secondary metabolite encoding gene clusters, and effectors in *A. brassicae*, and 4) carry out a comparative analysis of the genomes sequenced in this study with some of the previously sequenced *Alternaria spp.* genomes to gain insights into their pathogenic lifestyles.

## Materials and Methods

### 1. Genome Sequencing and assembly

Two isolates of *A. alternata* which were found to be co-infecting *B. juncea* leaves along with *A. brassicae* in our experimental field station at Delhi, India (PN1 and PN2) were isolated and purified by single spore culture. High molecular-weight genomic DNA was extracted from mycelia of 5-day old cultures of *A. alternata* isolates using a method described earlier^10^. 2 μg of the high molecular-weight genomic DNA was used for Nanopore library preparation using LSK-108 ONT ligation protocol. The libraries were then run on R9.4 SpotON MinION flowcells for 24 hours. Live base calling was enabled for all the runs. The MinION runs produced 4,14,210 and 2,68,910 reads amounting to 2.36 GB and 1.98 GB of data for *A. alternata* PN1 and PN2, respectively. The genomes were assembled *de novo* using the Canu assembler (version 1.6)^14^. Nanopolish was used to compute an improved consensus sequence using the signal-level raw data for the assemblies.

### 2. mRNA sequencing and transcript reconstruction

Total RNA was isolated from 15-day old fungal mycelia of *A. brassicae*, grown on Potato Dextrose Agar (PDA) and Radish Root Sucrose Agar (RRSA), using Qiagen RNeasy Plant Mini Kit following the manufacturer’s instructions. 100 bp sequencing libraries were prepared using the TruSeq Stranded mRNA Library Prep Kit (Illumina). Paired-end sequencing was carried out using Illumina SBS v4 chemistry on an Illumina Hiseq 2500. The Hiseq run generated 50 million PE reads per sample. The raw reads were trimmed using Trimmomatic, and the trimmed reads were then mapped to the *de novo* genome assembly using STAR (version 2.5.0)^15^. Transcripts were reconstructed using Cufflinks (version 2.2.1)^16^ and likely coding regions were identified using TransDecoder (version 5.3.0)^17^.

### 3. Gene prediction and annotation

RepeatModeler (version 1.0.11) was used for *de novo* repeat family identification. The *de novo* identified repeat library was used for masking the genome using RepeatMasker (version 4.0.7). The repeat-masked genome was used for gene predictions. For gene prediction, multiple lines of gene evidence were integrated using EVM (EvidenceModeler)^18^. Two *ab initio* gene callers were used viz. AUGUSTUS^19^ and GeneMark-HMM-ES^20^. GeneMark-HMM-ES was self-trained on the repeat-masked genome whereas AUGUSTUS was trained on the genome and cDNA hints from *A. alternata.* RNA-Seq evidence in the form of coding regions identified by TransDecoder was also used. Additionally, homology-based gene prediction was carried out using GeMoMa^21^ with protein-coding genes of *A. longipes, A. arborescens* and *A. alternata*.

Each of these lines of evidence was presented to EVM as separate tracks. In EVM, weights were assigned to each evidence as follows: AUGUSTUS 1, GeneMark-HMM-ES 1, GeMoMa 1, and RNA-Seq evidence 5. The genes predicted by EVM were used for all the downstream analyses. Genes were then annotated using BLAST against UniProt, SWISS-PROT, CAZy, MEROPS, and PHI-BASE. The fungal version of antiSMASH (version 4.0)^22^ was used to identify secondary metabolite gene clusters in the genomes. Candidate effector proteins were predicted using the following pipeline: a) SignalP (version 4.1)^23^ and Phobius to identify secreted proteins, b) TMHMM (version 2.0)^24^ to remove proteins with transmembrane domains, c) predGPI to filter out proteins that harbored a GPI membrane-anchoring domain, and d) EffectorP (version 2.0)^25^ to predict potential effectors from the above-filtered protein set.

### 4. Orthology and whole-genome phylogenetic analysis

The genomes of 16 *Alternaria* species (Supplementary Table 2) were included in the analysis with *Stemphylium lycopersici* as an outgroup. The proteomes of the fungi were clustered using the Orthofinder (version 2.2.6)^26^ pipeline with default parameters. The clusters were further analysed with Mirlo (https://github.com/mthon/mirlo) to identify phylogenetically informative single copy gene families. These families were then concatenated into one large alignment and used for phylogenetic analysis. A phylogenetic tree was constructed using Bayesian MCMC analysis based on the concatenated alignment under the WAG+I evolutionary model and the gamma distribution calculated using four rate categories and homogenous rates across the tree. Orthologs were also identified for 13 other pathogenic *Dothideomycetes* (Supplementary Table 2) for comparative analyses using the same pipeline as above.

### 5. Relationship of TEs and repeat-rich regions to genes and gene clusters

The overlap of repeat-rich regions and transposable elements (TEs) with i) genes encoding secreted proteins, ii) effectors and iii) secondary metabolite gene clusters were analysed using the regioneR^27^ package in R. The analysis involved comparison of the overlap of each the above gene sets with transposable elements and repeat-rich regions with a random set of genes selected from the genome. A distribution of means was established by running 10,000 permutation tests, which was then used to calculate a p-value.

## Results and discussion

### 1. Genomic features of *A. brassicae* and two other co-infecting *A. alternata* isolates

We sequenced the genomes of two isolates of *A. alternata* (PN1 and PN2) that were co-infecting *B. juncea* with *A. brassicae*. The *A. brassicae* assembly has been previously described^10^. Briefly, the assembly consisted of nine complete chromosomes and one chromosome with telomeric repeats missing at one of the ends. Apart from these chromosomes, there were six contigs of which one of them was ∼1 Mb in size, which may together constitute a dispensable chromosome. The N50 of the *A. brassicae* assembly was 2.98 Mb (Table 1). The two isolates co-infecting *B. juncea* were identified to be *A. alternata* based on their ITS and GAPDH sequences. The *A. alternata* assemblies Aat_PN1 and Aat_PN2 consisted of 14 contigs totalling to 33.77 Mb, and 15 contigs totalling to 33.53 Mb, respectively (Table 1). Six contigs in each of the two assemblies contained telomeric repeats on both ends and therefore, are most likely to represent full chromosomal molecules. Four other contigs in both the assemblies contained telomeric repeats on one end but were of similar size of full chromosome molecules as described in *A. solani*^*13*^. Therefore, the genome assemblies for *A. alternata* isolates represented ten nearly complete chromosomes each. Whole genome alignments with related *Alternaria spp.* showed an overall synteny between the genomes with minor rearrangements. Additionally, mitochondrial sequences were also obtained from the sequencing data for the two isolates of *A. alternata*. The mitochondrial genomes of the *A. alternata* strains were approximately 49,783 bp and 50,765 bp in size respectively and showed high similarity with the previously published mitochondrial genome of *A. alternata*^28^.

**Table 1:** Assembly statistics of the six near-complete *Alternaria* genome sequences.

|  | *A. brassicae*<br>J3<sup>10</sup> | *A. alternata*<br>PN1 | *A. alternata*<br>PN2 | *A. solani*<br>altNL03003<sup>13</sup> | *A. brassicicola*<br>abra43<sup>11</sup> | *A. alternata*<br>ATCC34957<sup>12</sup> |
| --- | --- | --- | --- | --- | --- | --- |
| **Assembly size (Mb)** | 34.14 | 33.77 | 33.53 | 32.78 | 31.04 | 33.48 |
| **No. of contigs** | 17 | 14 | 15 | 10 | 29 | 27 |
| **No. of contigs**<br>**( >10000 bp)** | 17 | 13 | 15 | 10 | 29 | 25 |
| **Largest contig (Mb)** | 7.1 | 6.86 | 6.76 | 6.94 | 3.3 | 3.96 |
| **N50** | 2.98 | 3.09 | 3.1 | 2.87 | 2.1 | 2.83 |
| **GC (%)** | 50.7 | 50.98 | 50.95 | 51.32 | 50.85 | 50.95 |
| **Repeat content (%)** | 9.33 | 2.43 | 2.64 | 5.71 | 9.3 | 2.71 |
| **Predicted genes** | 11593 | 11495 | 11387 | 11804 | 10261 | 12500 |

**Table 2:** Comparison of characteristics of Core chromosomes and dispensable chromosome of *A. brassicae*.

| Characteristic | Core chromosomes | DC largest contig (ABRSC11) | DC contigs (all) |
| --- | --- | --- | --- |
| Total length (bp) | 32140555 | 997589 | 1809659 |
| G+C (%) | 50.85 | 50.42 | 47 |
| Number of protein-coding genes | 11216 | 238 | 377 |
| Proportion of genes by length (%) | 52.48 | 30.86 | 30.05 |
| Number of Transposable element (TE) copies | 1454 | 181 | 313 |
| Proportion of TEs by length (%) | 5.78 | 21.45 | 20.89 |
| Proportion of secreted protein genes (%) | 10.09 | 9.66 | 9.81 |
| Proportion of effector genes (%) | 1.69 | 2.52 | 2.39 |

Gene prediction following repeat masking resulted in the identification of 11593, 11495, and 11387 genes in the *A. brassicae, A. alternata* PN1, and PN2 genome assemblies, respectively. This was comparable to the gene numbers estimated in other *Alternaria spp.* (Table 1). BUSCO analysis showed that the gene models predicted in the three genomes covered 98 % of the single copy conserved fungal genes indicating near-completeness of the assemblies. The predicted genes were comprehensively annotated using a combination of databases as described in the Methods section (Figure 1; Supplementary Table 1). In addition to the three genomes, we also predicted genes *de novo* in the genome assemblies of three other *Alternaria* species which were sequenced using long-read technologies viz. *A. brassicicola* (abra43)^11^, *A. alternata* (ATCC34957)^12^, and *A. solani* (altNL03003)^13^ (Table 1). These six genomes and their gene predictions were used for the comparative analyses of secondary metabolite encoding gene clusters and effector-coding genes.

**Figure 1:**
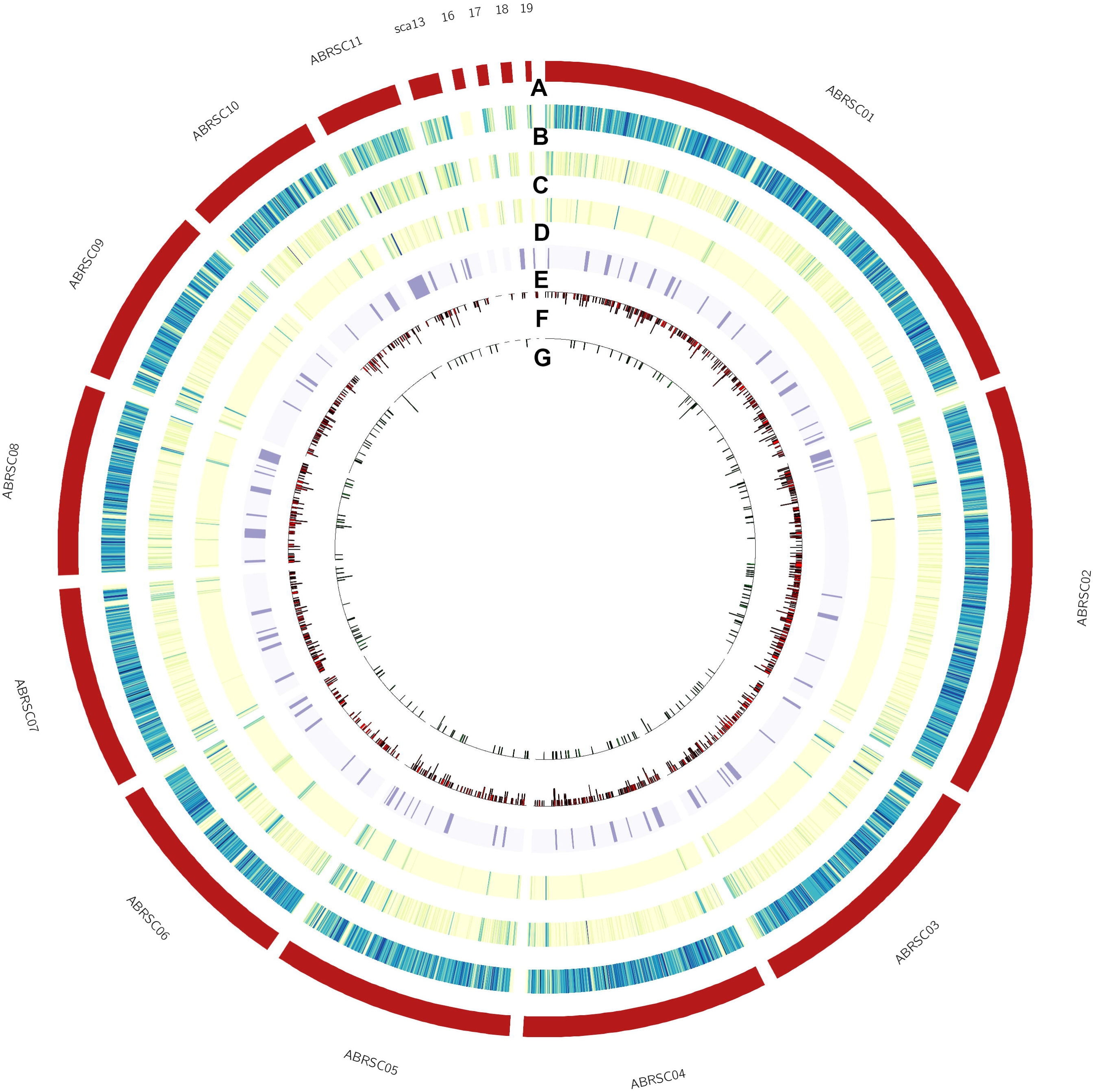
Summary of *A. brassicae* genome, (From outer to inner circular tracks) A) pseudochromosomes/scaffolds, B) Protein-coding genes, B) Repeat elements, D) Transposable Elements (DNA and LTR), E) predicted secondary metabolite clusters, F) Secreted proteins, G) predicted effectors.

### 2. Phylogenomic analysis assigns a separate clade for the Brassica-infecting *A. brassicae* and *A. brassicicola* within the *Alternaria* genus

In order to accurately reconstruct the divergence and relationship between *A. brassicae*, the two *A. alternata* isolates (PN1 and PN2), and the other *Alternaria* species, we conducted phylogenomic analyses using 29 single copy orthologs that had the highest phylogenetic signal as calculated by the program Mirlo. Selection of genes with higher phylogenetic signals leads to phylogenies that are more congruent with the species tree^29^. The resulting phylogeny showed that the large-spored *Alternaria* and small-spored *Alternaria* species clustered separately into two different clades (Figure 2). The *A. alternata* isolates PN1 and PN2 co-infecting Brassicas clustered together with the *A. alternata* isolated from sorghum (ATCC34957). Interestingly, the two major pathogens of the Brassicas viz. *A. brassicae* and *A. brassicicola* clustered separately from all the other *Alternaria* species, possibly indicating a very different evolutionary trajectory based on the host preferences of these two species.

**Figure 2:**
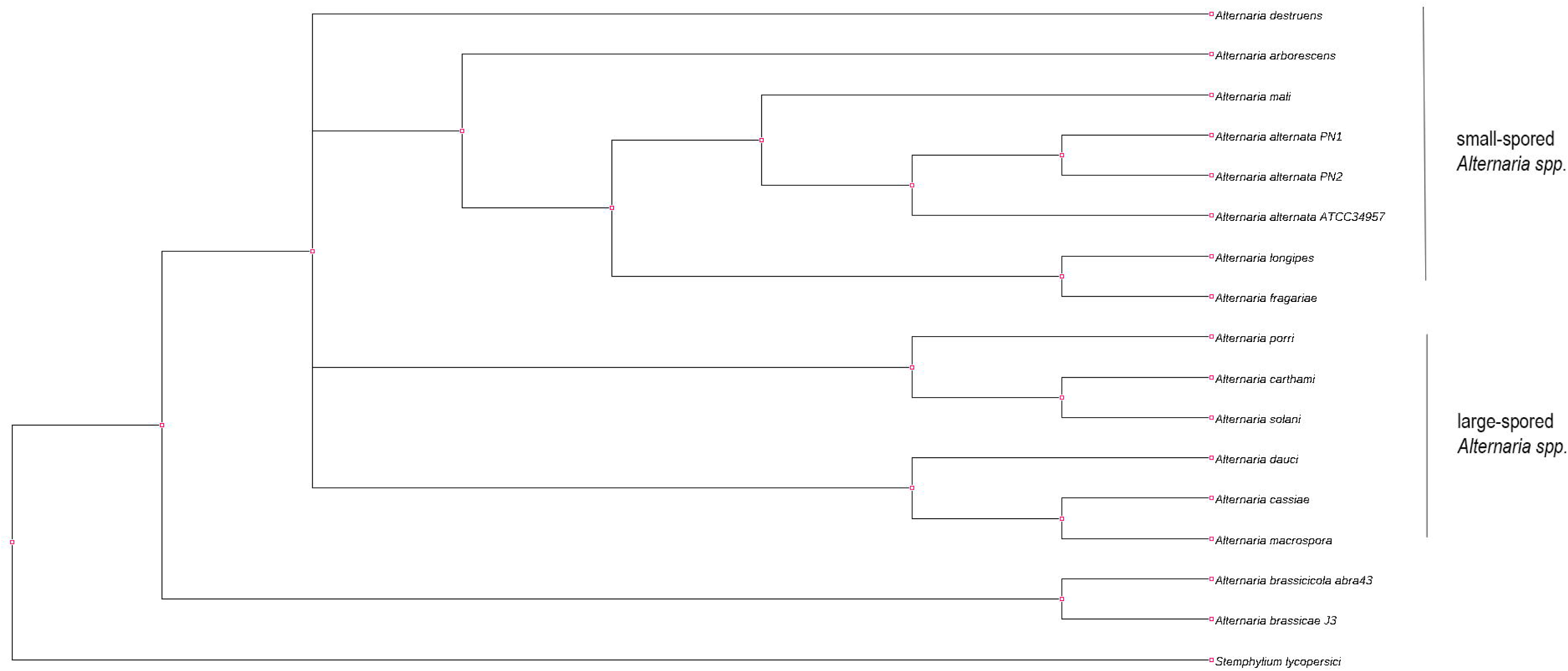
Phylogenetic tree of *Alternaria* species with *S. lycopersici* as an outgroup. The tree was constructed using 29 single copy orthologs, which had the highest phylogenetic signal as calculated in Mirlo.

### 3. Comparative analyses of *A. alternata* species isolated from different hosts

We compared the genomes of *A. alternata* PN1 and PN2 (isolated from *B. juncea*) to that of *A. alternata* ATCC34957 (isolated from sorghum) to identify any differences in their genomic content that might allow these to infect two very different species. Whole-genome alignments of *A. alternata* PN1 and PN2 to that of *A. alternata* ATCC34957 revealed very high levels of synteny and the absence of any species-specific regions. Notably, all the three species did not contain any dispensable chromosomes which may confer pathogenicity, as has been reported for *A. alternata* isolates infecting many of the fruit crops such as citrus, pear, and apple^31,32,45^. The gene repertoires of the three species also consisted of similar number and type of effectors, CAZymes, and secondary metabolite clusters (Table 1). These results suggest that these isolates of *A. alternata* may be opportunistic secondary pathogens.

### 4. An abundance of repeat-rich regions and transposable elements in *A. brassicae*

Filamentous plant pathogens tend to have a distinct genome architecture with higher repeat content. Many plant pathogens are known to have a bipartite genome architecture or what is generally known as a two-speed genome, in which the gene-sparse repeat-rich region provides the raw material for adaptive evolution^30^. Repeat content estimation and masking using RepeatModeler and RepeatMasker revealed that the *A. brassicae* genome consisted of ∼9.33% repeats as compared to 2.43% and 2.64% repeats in the *A. alternata* genomes. The *A. brassicae* genome harbors the highest repeat content (∼9.33%) among all the *Alternaria* species sequenced till date. However, the genome does not exhibit any bipartite genome architecture, as reported in some phytopathogens^30^. Our analysis showed that the repeat content differs significantly between the *A. alternata* isolates and the other pathogenic *Alternaria* species. The pathogenic *Alternaria* species especially *A. brassicae* and *A. brassicicola* had a considerably larger repertoire of LTR/Gypsy and LTR/Copia elements (> 8X) in comparison to the other *A. alternata* isolates (pathogenic and non-pathogenic) (Figure 3). The *A. brassicae* and *A. brassicicola* genomes also had an overrepresentation of DNA transposons, which amounted to ∼5% of the genome, as compared to <1% in the other *Alternaria* species (Figure 3).

**Figure 3:**
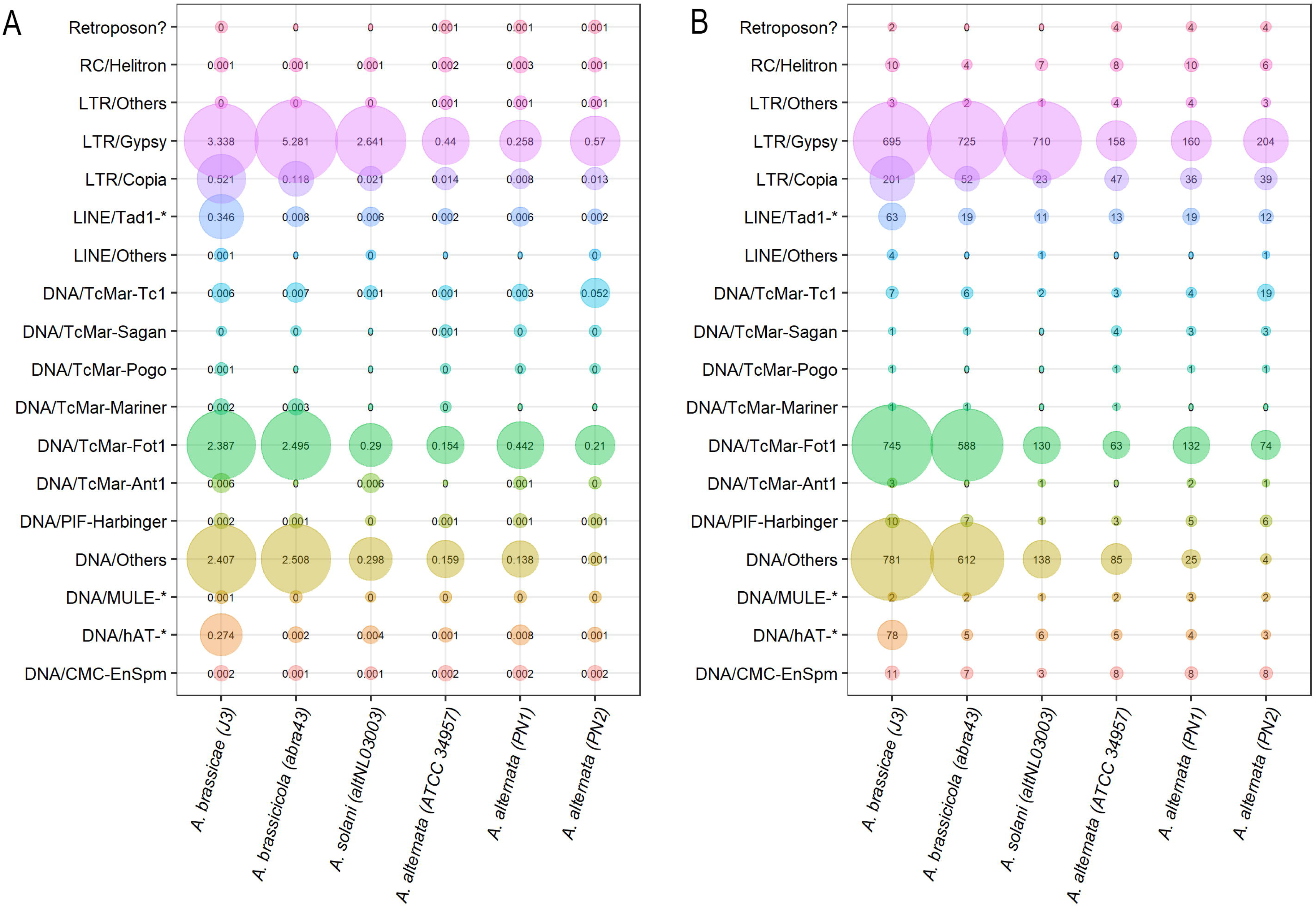
Comparison of repeat content in six *Alternaria* species. The size of the bubbles corresponds to the A) percentage of transposable elements (TEs) in the genome, B) copy number of the TE in the genome.

This proliferation of repetitive DNA and subsequent evolution of genes overlapping these regions may be the key to evolutionary success wherein these pathogens have managed to persist over generations of co-evolutionary conflict with their hosts. Proximity to TEs potentially exposes the genes to RIP-mutations and therefore accelerated evolution. Ectopic recombination between similar TEs may also result in new combinations of genes and thereby increase the diversity of proteins or metabolites.

### 5. Presence of a dispensable chromosome in the large-spored *A. brassicae*

Lineage-specific (LS) chromosomes or dispensable chromosomes (DC) have been reported from several phytopathogenic species including *A. alternata.* DCs in *A. alternata* are known to confer virulence and host-specificity to the isolate. The whole-genome alignments of *A. brassicae* with other *Alternaria* spp. revealed that a contig of approx. 1 Mb along with other smaller contigs (66-366 kb) was specific to *A. brassicae* and did not show synteny to any region in the other *Alternaria spp.* However, partial synteny was observed when the contig was aligned to the sequences of other dispensable chromosomes reported in *Alternaria spp*.^31,32^. This led us to hypothesize that these contigs together may represent a DC of *A. brassicae*. To confirm this, we searched the contigs for the presence of AaMSAS and ALT1genes, which are known marker genes for dispensable chromosomes in *Alternaria spp.*^4^. We found two copies of the AaMSAS gene as part of two secondary metabolite biosynthetic clusters on the 1 Mb contig. However, we did not find any homolog of the ALT1 gene. Additionally, the repeat content of the contigs (ABRSC11, scaffold 13, 17, 18, and 19) was compared to the whole genome. The gene content of the lineage-specific contigs was significantly lower than that of the core chromosomes (Table 2). Conversely, the DC contigs were highly enriched in TE content as compared to the core chromosomes (Table 2). Although, the DC was not enriched with genes encoding secreted proteins, the proportion of secreted effector genes was 30% higher as compared to the core chromosomes. All the above evidence point to the fact that *A. brassicae* may indeed harbour a DC. DCs in *Alternaria spp.* have been reported so far from only the small-spored *Alternaria spp.* and no large-spored *Alternaria* species have been known to harbour DCs. It remains to be seen whether the DC contributes to virulence of *A. brassicae*. Future studies would involve the characterization of the dispensable chromosome in *A. brassicae* and correlating its presence to the pathogenicity of different isolates.

### 6. Orthology analysis reveals species-specific genes with putative roles in virulence

Differences in gene content and diversity within genes contribute to adaptation, growth, and pathogenicity. In order to catalogue the differences in the gene content within the *Alternaria* genus and the *Dothideomycetes*, we carried out an orthology analysis on the combined set of 360216 proteins from 30 different species (including 16 *Alternaria* species) belonging to *Dothideomycetes* (Supplementary Table 2) out of which 3,45,321 proteins could be assigned to an orthogroup. We identified 460 *A. brassicae* specific genes which were present in *A. brassicae* but absent in all other *Alternaria* species (Supplementary Table 3). These species-specific genes included 35 secreted protein coding genes out of which 11 were predicted to be effectors. Additionally, 20 of these species-specific genes were present on the DC. A large number of these proteins belonged to the category of uncharacterised proteins with no known function. In order to test whether these species-specific genes are the result of adaptive evolution taking place in the repeat-rich regions of the genome, we carried out a permutation test to compare the overlap of repeat-rich regions and transposable elements with a random gene set against the overlap of these species-specific genes. We found that these species-specific genes overlapped significantly with repeat-rich regions (P-value: 9.99e-05; Z-score: −4.825) and transposable elements (P-value: 0.0460; Z-score: 2.539) in the genome.

### 7. Synteny analysis reveals the genetic basis of the exclusivity of Destruxin B production by *A. brassicae* within the *Alternaria* genus

The genera of *Alternaria* and *Cochliobolus* are known to be the major producers of host-specific secondary metabolite toxins. *Alternaria spp.* especially are known for the production of chemically diverse secondary metabolites, which include the host-specific toxins (HSTs) and non-HSTs. These secondary metabolites are usually generated by non-ribosomal peptide synthases (NRPS) and polyketide synthases (PKS). We identified five NRPS type SM gene clusters, 12 PKS type gene clusters and seven terpene-like gene clusters in *A. brassicae* (Supplementary Table 4). Out of the five NRPS clusters, we could identify three clusters which produce known secondary metabolites viz. Destruxin B, HC-toxin and dimethylcoprogen (siderophore).

Destruxin B represents a class of cyclic depsipeptides that is known to be one of the key pathogenicity factors of *A. brassicae* and has been reported to a host-specific toxin of *A. brassicae*^5^. Destruxin B has not been reported to be produced by any of the other *Alternaria* species. Here we report for the first time the biosynthetic gene cluster responsible for Destruxin B production in *A. brassicae*. The cluster consists of 10 genes, including the major biosynthetic enzyme encoded by an NRPS gene (DtxS1) and the rate-limiting enzyme, DtxS3 (aldo-keto reductase) (Supplementary Table 4). Interestingly, synteny analysis of this cluster among the six *Alternaria* species showed that both these genes were not present in any of the other *Alternaria* spp. although the overall synteny of the cluster was maintained in all of these species (Figure 4). The absence of the key genes coding for the enzymes DtxS1 and DtxS3 in the Destruxin B cluster in the other *Alternaria* species explains the absence of Destruxin B in those species.

**Figure 4:**
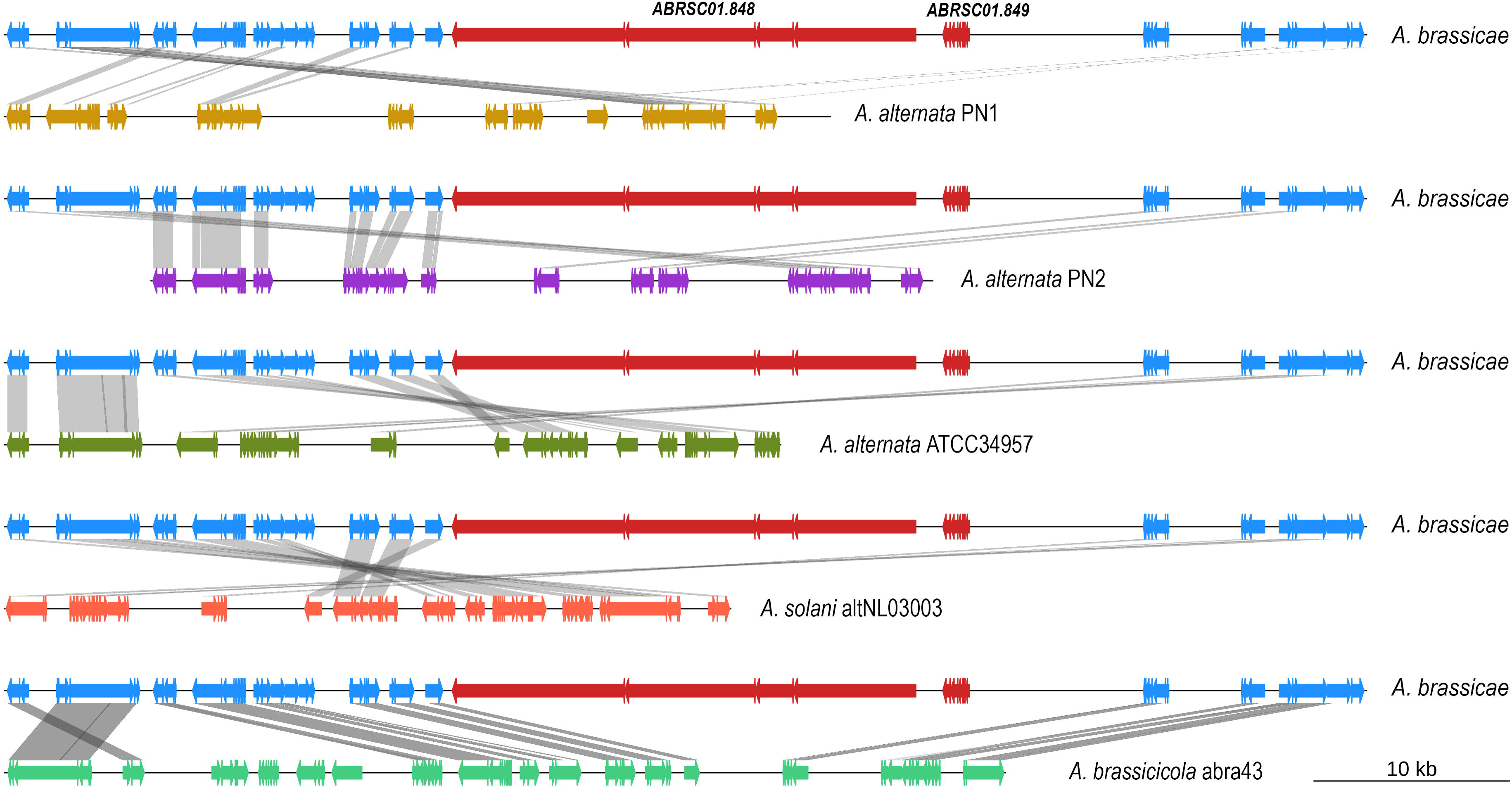
Synteny of the Destruxin B cluster in the six *Alternaria* species. DtxS1 and DtxS3 are marked in red (labelled with respective gene IDs) and are absent from all the other *Alternaria* species.

Destruxin B has been reported from the entomopathogen *Metarhizium robertsii*^33^, and *Ophiosphaerella herpotricha*^34^, the cause of spring dead spot of bermudagrass, apart from *A. brassicae*. Wang et al.^35^ described the secondary metabolite biosynthetic cluster responsible for Destruxin production in *M. robertsii*. The evolutionary history of Destruxin B production within the Metarhizhium genus closely resembled that of *Alternaria*. The specialist pathogens such as *M. acridum* lack the two key enzymes and hence do not produce Destruxins. However, *M. robertsii*, a generalist with a wider host range produces Destruxins^35^. It was therefore hypothesized that Destruxins might be responsible for the establishment of the extended host range of *M. roberstii*^35^. A similar hypothesis may also be true in the case of *A. brassicae*, which has a broad host range and can infect almost all the *Brassicaceae*. Further experiments to determine the host range of Destruxin null mutants of *A. brassicae* may help clarify the role of this important phytotoxin in extending the host range of *A. brassicae*.

We also identified a gene cluster responsible for dimethylcoprogen (siderophore) production in *A. brassicae*. Siderophores are iron□chelating compounds, used by fungi to acquire extracellular ferric iron and have been reported to be involved in fungal virulence^36^. The identification of the gene cluster responsible for siderophore synthesis would enable the study of siderophores and their role in pathogenicity in *A. brassicae*. Additionally, we identified an NRPS cluster, possibly coding for HC-toxin in one of the CDCs (scaffold 18) (Supplementary Table 4). HC-toxin is a known virulence determinant of the plant pathogen *Cochliobolus carbonum*, which infects maize genotypes that lack a functional copy of HM1, a carbonyl reductase that detoxifies the toxin^37^. A recent report showed that *A. jesenskae* also could produce HC-toxin, making it the only other fungus other than *C. carbonum* to produce the toxin^38^. The presence of the HC-toxin cluster in *A. brassicae* hints towards the possibility of many other species having the potential to produce HC-toxin outside the *Cochliobolus* genus. Additionally, a PKS type cluster consisting of 12 genes, responsible for melanin production was also identified (Supplementary Table 4). The melanin biosynthetic cluster has been described for *A. alternata* previously^39^. Also, the transcription factor Amr1, which induces melanin production, has been characterized in *A. brassicicola* and is known to suppress virulence^40^. The role of melanin in virulence is ambiguous. The melanisation of appressoria contributes to virulence in *Magnaporthe* and *Colletotrichum* species^41^. In the case of *Cochliobolus heterostrophus*, melanin deficient isolates were able to sustain virulence under lab conditions but lost their virulence in the field^42^. However, melanin production in *A. alternata* has been reported to play no role in its pathogenicity^43^.

The plant pathogens belonging to the genus of *Alternaria* seem to have a dynamic capacity to acquire new secondary metabolite potential to colonize new ecological niches. The most parsimonious explanation for this dynamic acquisition of secondary metabolite potential is horizontal gene transfer within the genus of *Alternaria* and possibly with other genera. There is extensive evidence in the literature that much of the HSTs of *Alternaria* are carried on the dispensable chromosomes and exchange of these chromosomes can broaden the host specificity^4,44,45^. Apart from horizontal gene transfer, rapid duplication, divergence and loss of the SM genes may also contribute to the pathogen evolving new metabolic capabilities. These processes of duplication and divergence may well be aided by the proximity of the secondary metabolite clusters to the repeat elements that makes them prone to RIP-mutations. Therefore, we tested whether the secondary metabolite clusters were also associated with repeat-rich regions. A permutation test was used to compare the overlap of repeat-rich regions with a random gene set against the overlap of secondary metabolite cluster genes. The secondary metabolite clusters significantly overlapped repeat-rich regions as compared to the random gene set (P-value: 0.0017; Z-score: −2.7963). Also, these clusters overlapped significantly with transposable elements among the repeat-rich regions (P-value: 0.0087; Z-score: 2.9871). This shows that both the mechanisms described above for the acquisition of new secondary metabolite potential may be possible in the case of *A. brassicae*. Population-scale analyses at the species and genus level may throw light on the prevalence of these mechanisms within the genus of *Alternaria*.

### 8. Distinct CAZyme profiles of *A. brassicae* and *A. brassicicola* within the *Alternaria* genus

CAZymes (Carbohydrate-Active enZymes) are proteins involved in the degradation, rearrangement, or synthesis of glycosidic bonds. Plant pathogens secrete a diverse range of CAZymes that breakdown the complex polysaccharides in the plant cell wall. They consist of five distinct classes *viz*. Glycoside hydrolases (GH), Glycosyltransferases (GT), Polysaccharide lyases (PL), Carbohydrate esterases (CE), and Carbohydrate-binding modules (CBM). We identified > 500 CAZymes in the six *Alternaria spp.* including *A. brassicae* (Supplementary Table 5). The CAZyme distribution of *A. brassicae* and *A. brassicicola* varied from those of the other *Alternaria spp.* thus forming a separate cluster (Figure 5). The number of auxiliary activity enzymes or the enzymes involved in plant cell wall degradation varied considerably between the different genera compared. Nearly 46% of the CAZymes in *A. brassicae* were secreted out of which ∼17% were predicted to be effectors. Eight of the CAZy families were exclusively found in the *Alternaria* genus but not in the *Cochliobolus* and *Zymoseptoria* genus whereas there was a complete absence of GH39 family in all the *Alternaria* species but present in the other two genera. The AA9 family (formerly GH61; copper-dependent lytic polysaccharide monooxygenases) is significantly enlarged in comparison to the other CAZy families in the *Alternaria* and *Cochliobolus* genera with each species containing > 20 copies of the gene. The copy numbers in the *Alternaria spp.* are much higher than the copy numbers reported for *Botrytis* and *Fusarium spp.*^46^. The AA9 family is involved in the degradation of cell-wall polysaccharides and are known to act on a range of polysaccharides including starch, xyloglucan, cellodextrins, and glucomannan. LPMOs have been hypothesized to have a dual role – directly cleaving the cell-wall polysaccharides, and acting as a ROS generator and thus contributing to the oxidative stress leading to necrosis in the plant tissues^47,48^. Strikingly, 11 of the 26 AA9 proteins present in *A. brassicae* are predicted to be secreted effectors. Characterisation of these CAZymes and their role in pathogenesis could be the subject of further studies.

**Figure 5:**
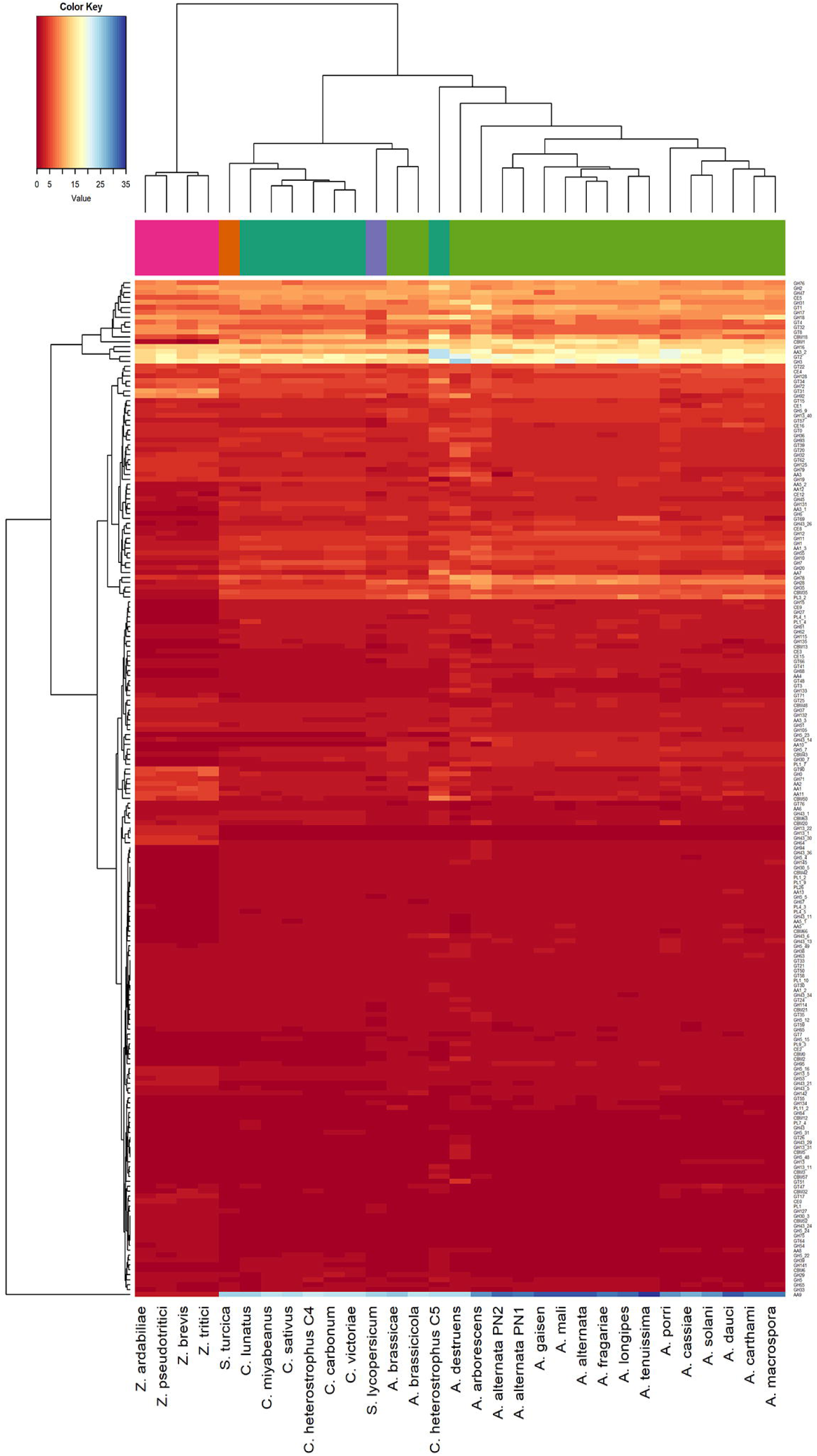
Heatmap of the CAZyme profiles in the 30 Dothideomycetes, which consists of the genera of *Alternaria, Cochliobolus*, and *Zymoseptoria*. Both the CAZyme families and the organisms were clustered hierarchically. The clustering of the organisms closely resembles that of the species phylogeny except for *A. brassicae* and *A. brassicicola*, which cluster separately from the other *Alternaria* species.

### 9. Overlapping effector profiles within the *Alternaria* genus

We predicted the effector repertoire of six *Alternaria* species (Table 1) including *A. brassicae* using the pipeline described in the methods section. Most of the small secreted proteins predicted to be effectors were uncharacterised/predicted proteins and thus may be novel species or genus-specific effectors. *A. brassicae* had the largest proportion of unknown/predicted/hypothetical proteins in the set of candidate effectors (Supplementary Table 6). We found that most of the effectors between the six species to be common and overlapping, suggesting that the broad mechanisms of pathogenesis involving proteinaceous effectors may be conserved within the genus. However, we found two copies of a beta/gamma-crystallin fold containing protein to be present only in *A. brassicae* and *A. brassicicola* and not in the other four *Alternaria* species. A further search through the nr database of NCBI revealed that this protein was completely absent in the *Alternaria* genus and the closest matches were found in other plant pathogens viz. *Macrophomina* and *Fusarium* species. However, no information is available as to its function or role in pathogenicity in any of the species.

We could also establish that some of the effectors in *A. brassicae* have the potential to evolve adaptively since they were also significantly associated with the repeat-rich regions of the genome (P-value: 0.0003; Z-score: −2.8778). Population-level analyses are therefore required to identify the effectors under positive selection, which could shed light on the evolution of pathogenicity in *A. brassicae*.

The effectors identified in this study reveal the wide range of proteins that may be involved in the pathogenesis of *A. brassicae*. 39 of these effectors were predicted to be CAZymes having various roles in the degradation of the cell wall and associated polysaccharides. The genome of *A. brassicae* contained two necrosis and ethylene-inducing peptide (NEP) proteins, which have been implicated in the pathogenesis of various pathogens including oomycetes and necrotrophs^49-51^. Hrip1 (Hypersensitive response inducing protein 1) from *A. alternata* has recently been shown to be recognized by MdNLR16 in a classical gene-for-gene manner, and deletion of Hrip1 from *A. alternata* enhances its virulence^52^. A Hrip1 homolog is also present in *A. brassicae*, but it is not predicted to be secreted outside the cell, although this needs to be verified experimentally. The presence of effectors which are recognized in a gene-for-gene manner opens up the possibility of identification of complementary R-genes in the host that can be utilized for developing resistant varieties or cultivars.

## Conclusions

*A. brassicae* has an enormous economic impact on the cultivated *Brassica* species worldwide, particularly the oleiferous types. Using the recently published high-quality genome assembly of *A. brassicae*, we annotated the genome and carried out comparative analyses of *A. brassicae* with other *Alternaria spp.* to discern unique features of *A. brassicae* vis-à-vis the other *Alternaria* species. We sequenced and annotated the genomes of two *A. alternata* isolates that were co-infecting *B. juncea.* The two *A. alternata* isolates had a gene content, effector repertoire, and CAZyme profiles that were very similar to that of an earlier sequenced *A. alternata* isolate (ATCC34957). This leads us to conclude that these isolates are opportunistic pathogens with a limited ability to cause infection on their own but would contribute overall to the disease outcome of a primary *A. brassicae* infection. Additionally, we show the presence of a dispensable chromosome in *A. brassicae*, a large-spored *Alternaria* species for the first time. The implications of a lineage-specific dispensable chromosome in *A. brassicae* towards pathogenesis remains to be unravelled. We also described the CAZyme profiles of nearly 30 *Dothideomycetes* and show that the CAZyme profiles of *A. brassicae* and *A. brassicicola* are different from the other *Alternaria* species. We also identified several important secondary metabolite gene clusters with putative roles in pathogenicity. The identification of the biosynthetic cluster responsible for Destruxin B in *A. brassicae* paves the way for reverse genetics studies to conclusively determine the contribution of Destruxin B towards the pathogenicity of *A. brassicae*. The repertoire of effectors identified in the six *Alternaria* species was largely overlapping. It may thus be hypothesised that host-specificity in the *Alternaria* species may be conferred by the combined action of proteinaceous effectors and the secondary metabolite toxins. Future studies would involve characterisation of the effectors and secondary metabolite clusters identified in this study and elucidating their role in pathogenesis.

## Availability

The genome assembly and associated raw data generated in this study have been deposited as National Center for Biotechnology Information BioProject PRJNA548052 and PRJNA548054.

## Acknowledgements

This work was supported by the grants from the Department of Biotechnology (DBT), Government of India, under the projects BT/IN/Indo-UK/CGAT/12/DP/2014-15 and BT/01/NDDB/UDSC/2016.

## Supplementary Tables

**Supplementary Table 1**: The complete annotated gene set of *A. brassicae* with gene coordinates, gene description and functional classification

**Supplementary Table 2**: List of *Dothideomycetes* used in the orthology analysis

**Supplementary Table 3**: List of 460 *A. brassicae* species-specific proteins with gene coordinates and description

**Supplementary Table 4**: List of predicted secondary metabolite gene clusters in *A. brassicae* along with their constituent genes, coordinates in the genome, and their description.

**Supplementary Table 5**: Comparison of the CAZyme profiles of the 30 *Dothideomycetes* including 16 *Alternaria* species

**Supplementary Table 6**: List of predicted effectors of *A. brassicae*

